# Synesthesia does not help to recover perceptual dominance following flash suppression

**DOI:** 10.1101/2019.12.12.873992

**Authors:** Diana Jimena Arias, Dave Saint-Amour

## Abstract

Grapheme-color synesthesia is a perceptual phenomenon that occurs when letters or numbers elicit an abnormal color sensation (e.g., printed black letters are perceived as colored graphemes). Grapheme-color synesthesia is typically reported following explicit presentation of graphemes. Very few studies have investigated color sensations in synesthesia in the absence of visual awareness. To address this issue, we took advantage of the dichoptic flash suppression paradigm to temporarily render a stimulus presented to one eye invisible. Synesthesic alphanumeric and non-synesthetic abstract stimuli were presented to 11 synesthete and 11 matched control participants in achromatic and chromatic experimental conditions. The test stimulus was first displayed to one eye and then masked following the sudden presentation of visual noise in the other eye. The time for an image to be perceived following the onset of the suppressive noise was calculated in each condition. Trials free of flash suppression but mimicking the perceptual suppression of the flash were also tested. Results showed that target detection by synesthetes was significantly better than by controls in the absence of flash suppression. However, no statistically significant difference was found between the groups when the test stimulus was interocularly suppressed, either for synesthetic or non-synesthetic stimuli. This study suggests that synesthesia can be associated with enhanced perception for overt recognition, but does not occur in the absence of visual awareness.

## Introduction

Synesthesia is a perceptual phenomenon in which a stimulus elicits an abnormal and concurrent sensation in the same or different sensory modality. Among the different types of synesthesia, grapheme-color synesthesia is relatively common and it consists of color perception evoked by grey scale alphanumeric images (letters and/or digits) [1]. Synesthetic associations are consistent over time [2–4] and are automatic or difficult to discard when elicited [3, 5–9].

Grapheme-color synesthesia has been most frequently investigated through the explicit presentation of synesthetic graphemes or digits, as shown with stroop-like tasks and visual search paradigms [5–10]. However, synesthesia remains much less studied when observers are unaware of a stimulus [for a review see 11]. Mattingley et al. [9] adapted a masking paradigm in order to test synesthetic stimuli under conditions preventing their visibility. They measured how long it takes for participants to name the color of a target mask that precedes the presentation of a letter prime, which evokes a synesthetic color, either congruent or incongruent with the color of the target mask. Congruence and incongruence trials were used to assess the synesthetic interference effect on color target naming. The letter prime was visible when presented for 500 ms and invisible when presented for 56 ms or less. The synesthetic interference effect was observed only when the prime was visible to participants, i.e., this effect disappeared when the letter did not access visual awareness. Furthermore, synesthete participants showed implicit priming effects similar to controls for non-inducing synesthetic stimuli; that is, the brief presentation of a semantic prime (e.g., an upper-case letter “A”) improved the subsequent recognition of a congruent target stimulus (e.g., lower-case letters “a”).

Synesthesia was also tested using the attentional blink paradigm in which two successive targets, T1 and T2, are presented in a sequence and separated by distractors (masks) [12]. In this paradigm, the second target T2 is rendered invisible when the presentation time between both targets does not exceed the attentional window, which is between 300 and 500 ms. Rich et al. [13] presented a synesthetic prime (T2) within and outside of the attentional window. A color probe was presented at the end of the attentional blink sequence. The prime elicited a synesthetic color that was either congruent or incongruent with the color probe, producing an interference effect in the color naming of the probe. Albeit modest, a reliable interference effect was found when the prime was visible. No interference effect was obtained when the prime was presented within the temporal window of the attentive blink. In another study using the same attentional blink design, however, some synesthetic participants (5 out of 10) were able to perceive synesthetic colors, even when the synesthetic primes fell within the attentional blink temporal window [14]. An interference effect in a color naming task was still noticed when participants were unable to overtly identify the synesthetic prime in the visual sequence. In line with this finding, it has been suggested in four grapheme-color synesthetes that conscious letter recognition is not required to perceive the color of hidden letters [15]. In this study, synesthete participants were able to identify unrecognizable letters (i.e., letters presented in mirror-reversed form or embedded in sagittal-plain words) much faster than non-synesthetes [15].

The aforementioned studies are not conclusive with regard to whether synesthesia can occur without visual awareness [9, 13–15]. To further investigative this question, we looked at the flash suppression paradigm, which has been used extensively to study visual processing in the absence of awareness [16, 17]. The flash suppression phenomenon is typically induced by two different monocular images presented asynchronously; an image is first presented to one eye for a few seconds (while a blank field is presented to the other eye), after which the second image is abruptly shown, i.e., flashed, to the other eye at the corresponding retinal points. Unlike the masking and attentional blink paradigms, flash suppression allows a stimulus to be rendered invisible temporarily, even though it remains physically present for the observer. Thus it allows manipulation of the onset of interocular suppression, i.e., the awareness of the suppressed stimulus, before binocular rivalry between the competing images occurs [16]. Unlike binocular rivalry, which involves alternation of perceptual dominance and complex neural dynamics at several brain levels [18], flash suppression offers better control of the monocular suppression [19]. Previous studies using flash suppression have shown that even if subjects are not aware of the presence of a stimulus in one eye, visual processing of that stimulus may still occur [16, 20–22].

In order to address whether synesthesia can occur in the absence of visual awareness, we used flash suppression to render the synesthetic stimulus invisible. By measuring the time the hidden stimulus takes to break suppression, it is possible to estimate the potential effect of synaesthesia on interocular suppression [20]. Importantly for the present study, it was previously shown in normal observers that the flash suppression of a colored Gabor grating is shorter than that of an achromatic Gabor grating [17]. Furthermore, some features of a stimulus, such as color, can break suppression more rapidly than other features, such as orientation [23]. Here we predicted that if synesthesia has the potential to occur when the participants are not aware of the synesthetic stimulus, these participants will exhibit a shorter duration of suppression in comparison to non-synesthetic stimuli and control participants.

## Materials and methods

### Participants

Eleven grapheme-color type synesthetes and 11 control participants were recruited to take part in this experiment. Synesthetes were matched to controls based on age (21-32 years old) and sex (4 men, 7 women). Grapheme-color associations in synesthetes were assessed qualitatively during a semi-structured interview and quantitatively using the grapheme-color consistency test from the online Synesthesia Battery, developed by Eagleman [24]. Synesthete participants were assessed twice using this battery, with a minimum lapse of two months between each testing session. Consistency test scores below 1.0 are indicators of synesthetic associations. Scores between 1.0 and 2.0 are not sufficiently conclusive to consider the presence of synesthetic associations, while scores higher than 2.0 rule out the possibility altogether. In our study, the average consistency scores for synesthetic participants between testing and re-testing were within the range of synesthetic associations (see Table 1). More precisely, the minimal average score for consistency was 0.43, while the maximum average score for consistency was 0.91. These values confirmed that the synesthetic associations reported by the participants were highly consistent over time. None of the participants had a history of neurological or psychiatric disorders, and reported normal or corrected-to-normal vision. Visual acuity was measured using the Snellen acuity chart and the contrast sensitivity FACT test (Stereo Optical Company Inc., Chicago, IL, United States). Stereoscopic vision was assessed using the Randot Test (Stereo Optical Co., Inc., Chicago, IL). All participants consented to participate in this study and received a financial compensation ($20 CAN). The experimental procedure conformed to the World Medical Association’s Declaration of Helsinki and was approved by the Research Ethics Committee of the Université du Québec à Montréal (FSH-2013-92).

**Table 1.** Demographic data, scores for the online synesthesia measures.

| Synesthete participant | Chronological age (years: months) | Sex | Consistency score | Projector/ Associator score |  | Alphanumeric stimulus |
| --- | --- | --- | --- | --- | --- | --- |
| 1 | 32:11 | F | 0.91 | -0.60 | Associator | 5 (red) |
| 2 | 32:06 | F | 0.73 | 0.09 | Projector | 5 (blue) |
| 3 | 23:11 | F | 0.43 | -0.17 | Associator | 5 (red) |
| 4 | 24:10 | F | 0.65 | -1.75 | Associator | 5 (red) |
| 5 | 22:03 | M | 0.49 | -1.92 | Associator | 5 (fuchsia) |
| 6 | 25:00 | F | 0.53 | -1.83 | Associator | 5 (orange) |
| 7 | 27:00 | M | 0.85 | -2.33 | Associator | 5 (red) |
| 8 | 21:04 | F | 0.80 | -1.42 | Associator | 5 (green) |
| 9 | 28:10 | F | 0.51 | -0.83 | Associator | 5 (green) |
| 10 | 31:07 | M | 0.73 | -2.75 | Associator | d (blue) |
| 11 | 23:07 | M | 0.42 | -0.57 | Associator | 5 (orange) |

Table 1 shows consistency and projector/associator scores from the synesthesia battery, and the selected synesthetic stimuli used for the current study. Negative scores for the projector/associator test mean that synesthetic associations are perceived in the “mind’s eye” (associator indicator) while positive scores indicate an “out of mind” synesthetic perception (projector indicator).

### Stimuli and design

Three types of stimulus were used: a synesthetic stimulus, a non-synesthetic stimulus, and a suppressor stimulus. The synesthetic stimulus was a number “5” for all participants except one (see Table 1), while the non-synesthetic stimulus was a symbol created from the trait features of the respective alphanumeric synesthetic (see Fig 1). The number “5” was chosen because it evokes a vivid synesthetic sensation of color in all participants. For synesthete participant #10, the synesthetic stimuli was replaced by a letter “d”, as this participant experienced no synesthesia with digits. The size of the stimuli was 1° x 0.6°. Stimuli were presented on a black square. The suppressor stimulus (1° x 1°) was a visual noise composed of random grains, ranging from black to white. These stimuli were presented side by side with at a distance of 3° from the central fixation point of the screen (see Fig 2).

**Fig 1.**
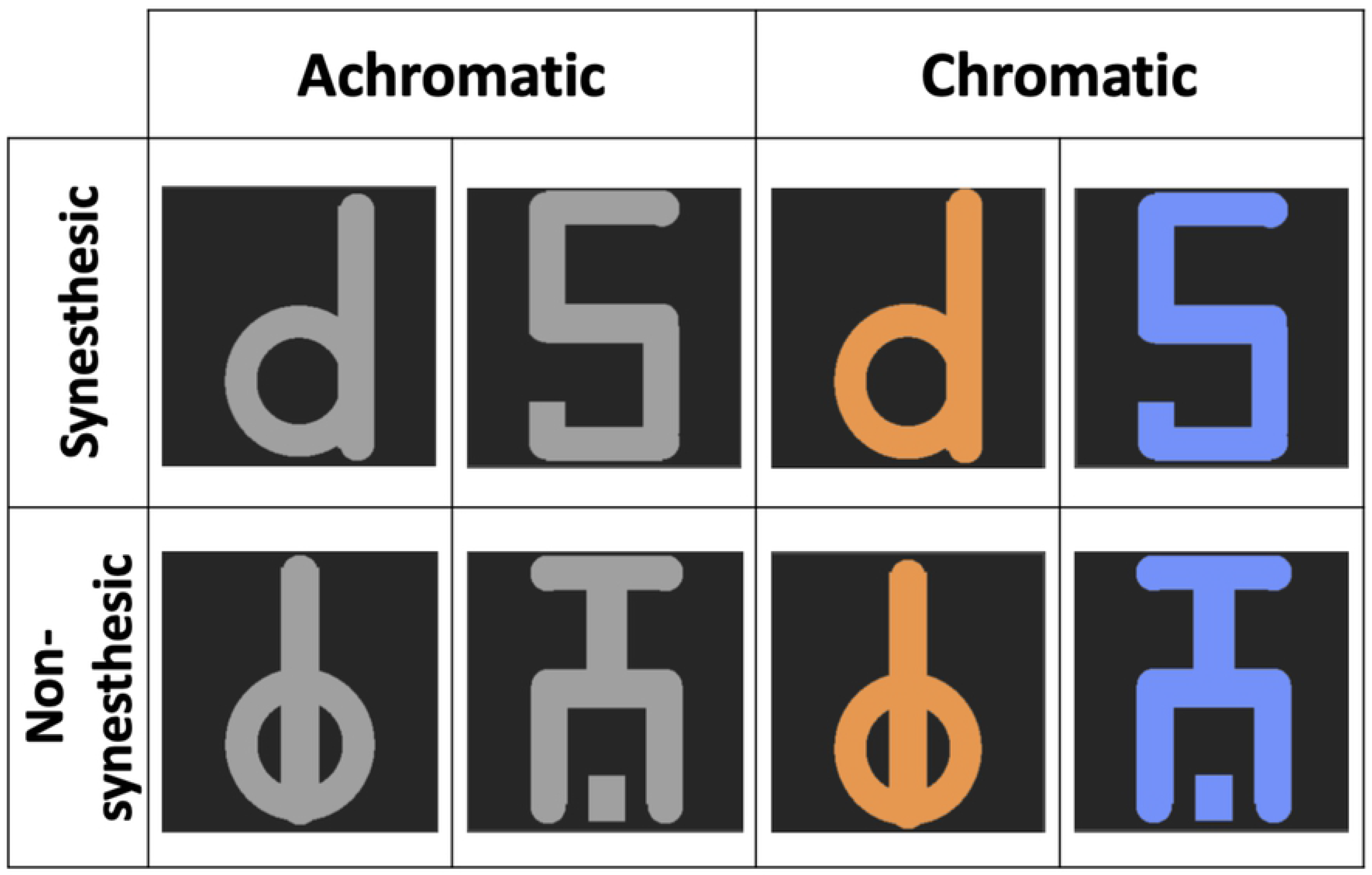
Stimuli. Synesthetic alphanumeric (first row) and non-synesthetic abstract stimuli (second row) were randomly presented.

Stimuli were generated and controlled with Psykinematix sofware, version 1.5 (KyberVision, Sendai, Japan). They were presented dichoptically using 3D glasses (head-mounted virtual-reality display model Z800 3D-Visor; eMagin Corp, Bellevue, WA) driven by a MAC G4 Desktop with an NVIDIA graphics card (GeForce 9400M, Santa Clara, CA). The resolution of each monocular, organic light emitting diode (OLED) screen was 800 by 600 pixels. In each OLED, the refresh rate was 60 Hz and the visual field was 32 by 23 degrees. The size of a pixel subtends an angle of 144 arc/sec (0.04 degrees).

Synesthetic and non-synesthetic stimuli, that is, alphanumeric and abstract stimuli, respectively, were displayed in two experimental conditions. In the achromatic condition, stimuli were presented in grey scale (RGB values = 160). In the chromatic condition, stimuli were displayed with the color that corresponded to the personal synesthetic perception of each synesthete. The RGB values of the images were then adjusted in order to reach physical equi-luminance with the achromatic stimuli (~ 49 cd/m2). All stimuli in all conditions were presented on a black background (0.01 cd/m2). The stimuli covered approximately 70% of the surface of the background frame. The contrast level was 50 %. Participants in the control group were tested with the same stimuli as their corresponding synesthete participant.

Four stimulus sets were generated according to the condition (achromatic and chromatic) and stimulus type (synesthetic and non-synesthetic): alphanumeric achromatic stimulus, achromatic abstract stimulus, alphanumeric chromatic stimulus, and abstract chromatic stimulus. Stimuli were displayed randomly in 3 blocks, each comprising 28 flash suppression trials. In the flash suppression trials, the presentation of stimuli was dichoptic (See Fig 2). A pair of suppressor stimuli (visual noise patches) was abruptly presented to the previously-unstimulated eye 3 to 4 seconds after the stimulus onset (time 0). As a result, the initial pair of target stimuli disappeared from awareness. Thus, the presentation of the visual noise was perceived by participants as a “flash,” which masked the test stimulus, even though it was physically present on the screen.

**Fig 2.**
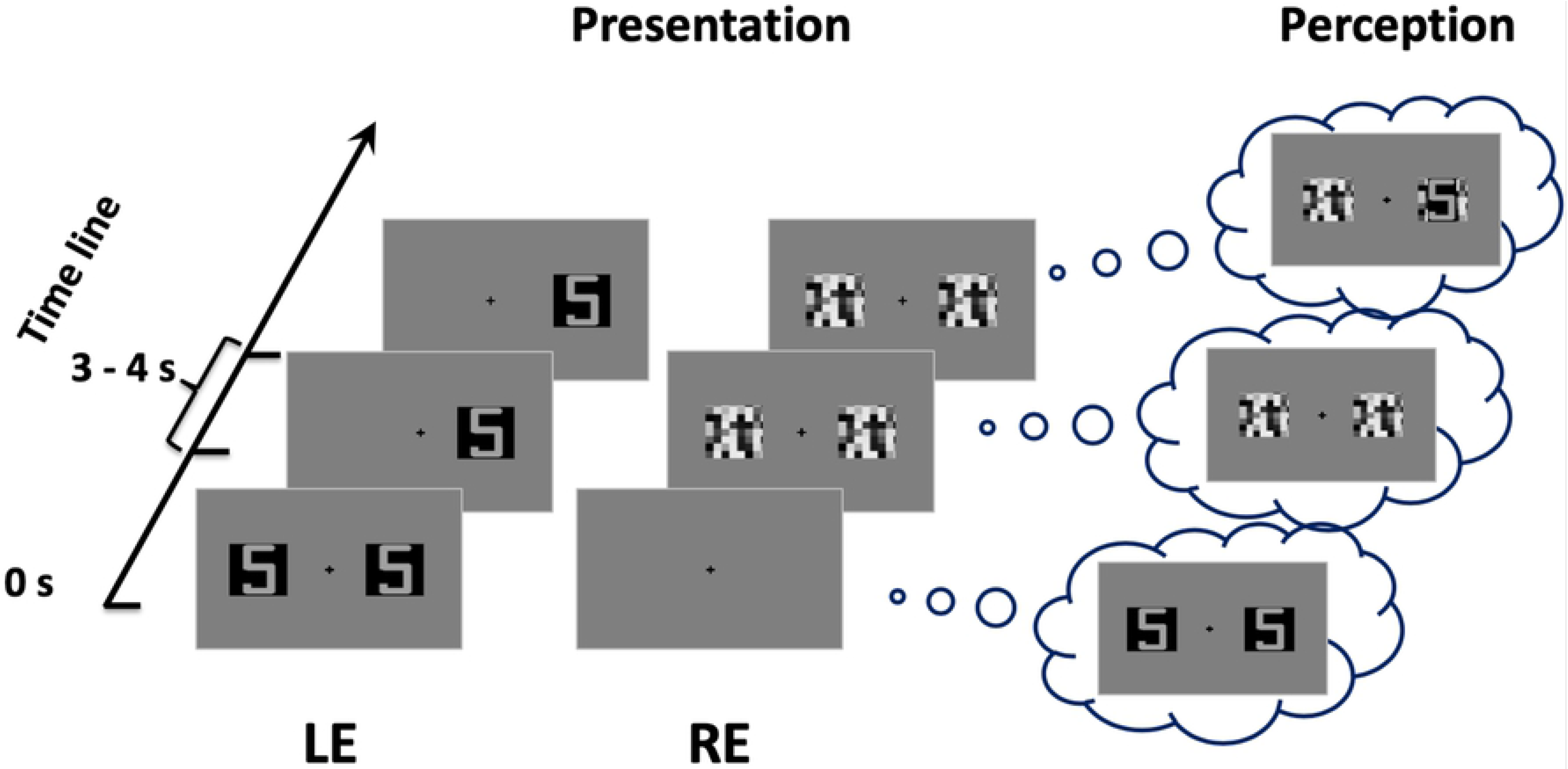
Example of a flash suppression trial. Following the noise patch stimulation in one eye, the initial pair of images in the opposite eye disappeared and one of the two images (either on the right or left side) reappeared after some time.

In addition to the flash suppression trials, “non-flash-suppression” trials (12 trials per block) were embedded in the task to mimic a stimulus presentation similar to the experimental trials. In the non-flash-suppression trials (See Fig 3), stimuli were presented in such a way that flash suppression did not occur; the initial stimuli displayed to one eye disappeared after 3-4 seconds. At the same time the two noise patches were presented to the other eye. After a while, the stimulus target reappeared slowly in one eye (fade-in, from 0 to 50% contrast) while the corresponding noise patch in the other eye disappeared slowly (fade-out, from 50 to 0%). The flash suppression (Fig 2) and the non-flash-suppression trials (Fig 3) were randomly presented in each testing block. In addition, stimulus presentation was counterbalanced between the left and the right eyes.

**Fig 3.**
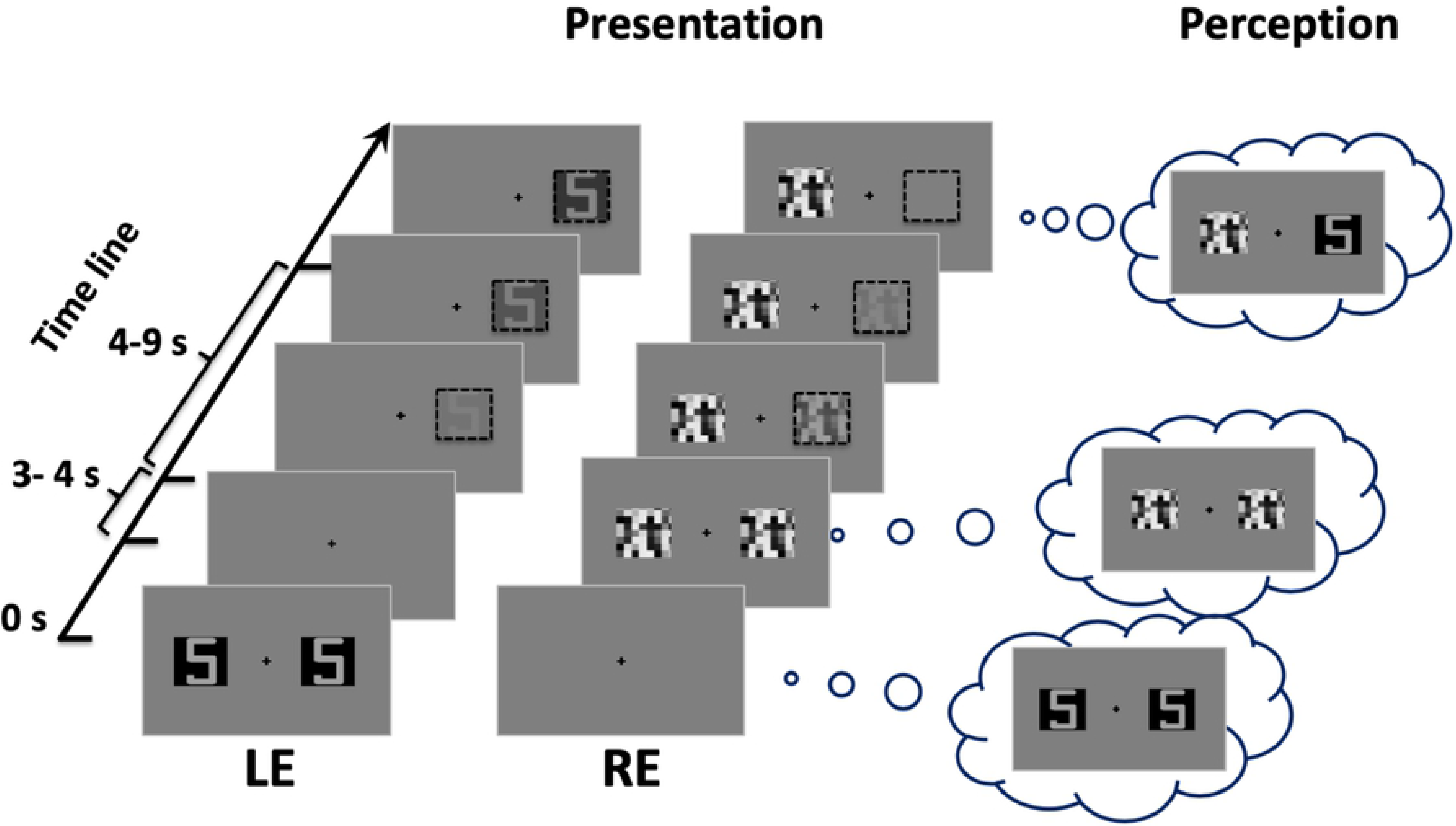
Example of a non-flash-suppression trial. The stimulus presentation was controlled (left side) to mimic the subjective perception (right side) so that one of the two images initially presented reappeared after being suppressed by the sudden presentation of a flash. Dashed-line squares (on the left) illustrate the fade-in/fade-out interplay of the images that was used to mimic the breaking of the flash suppression.

### Procedure

Participants were comfortably seated in a dimly lit room. They were instructed to adjust the alignment of the 3D glasses by moving the lenses sideways and to adjust their proximity. Eight practice trials were performed. The participants were instructed to maintain their gaze on the central fixation dot while the images were presented.

The task method consisted of a spatial two-alternative forced choice. The participants were instructed to press the left or the right arrow key when one stimulus re-appeared in its entirety, either on the left or the right side to the screen. The time required for a participant to report the reappearance of the hidden stimulus was calculated as the duration of suppression in the flash suppression trials. For the non-flash-suppression trials, the time required to detect the fake suppressed stimulus was measured. Both types of trials (flash suppression and non-flash-suppression) ended when participants responded, or after 15 seconds. Each trial was launched by pressing the “enter” key. Participants were allowed to take breaks between blocks if they desired.

### Data analyses

Intra- and inter-group differences were assessed using analyses of variance (ANOVA) with condition (achromatic and chromatic), stimulus (alphanumeric and abstract stimuli), and group (synesthetes and controls) as the main factors. Separate ANOVA were conducted for the flash suppression and the non-flash-suppression trials. For the flash suppression trials, the duration of suppression was calculated by subtracting the onset of the suppressor stimulus presentation (noisy patches) from the participants’ reaction time. A lower value meant a faster time for the target to reach visual awareness. For the non-flash-suppression trials, the time required to detect the fake suppressed stimulus was calculated from the fade-in onset of the stimulus target to the reaction time of the participant. A lower value meant that such stimulus was rapidly detected. In all analyses, the *p* values were set to be significant at an α level of < 0.05. Bonferroni corrections were applied to detect the significance in post-hoc pairwise comparisons.

## Results

One synesthete (participant 6) consistently reported longer RTs for the non-flash-suppression trials, significantly lower than those obtained from the rest of the participants (*Z-scores* ≤ – 3.5). This participant was thus excluded from the main data analysis.

The ANOVA conducted on perceptual flash suppression trials showed no main effect of group [*F*_(1,19)_ = 0.523, *p* = 0.478] or interaction of group with the other factors: condition*group [*F*_(1,19)_ = 0.280, *p* = 0.603], stimulus*group [*F*_(1,19)_ = 0.716, *p* = 0.408] and the condition*stimulus*group [*F*_(1,19)_ = 0.001, *p* = 0.975]. However, a main effect of condition [*F*_(1,19)_ = 63.833, *p* < 0.001, η^2^ = 0.771], stimulus [*F*_(1,19)_ = 21.591, *p* < 0.001, η^2^ = 0.532] as well as the interaction condition* stimulus [*F*_(1,19)_ = 36.989, *p* < 0.001, η^2^ = 0.661] was found to be statistically significant. As is depicted in Fig 4, all participants exhibited a significantly shorter duration of suppression in the chromatic condition than in the achromatic condition, and this effect was stronger for alphanumeric stimuli.

**Fig 4.**
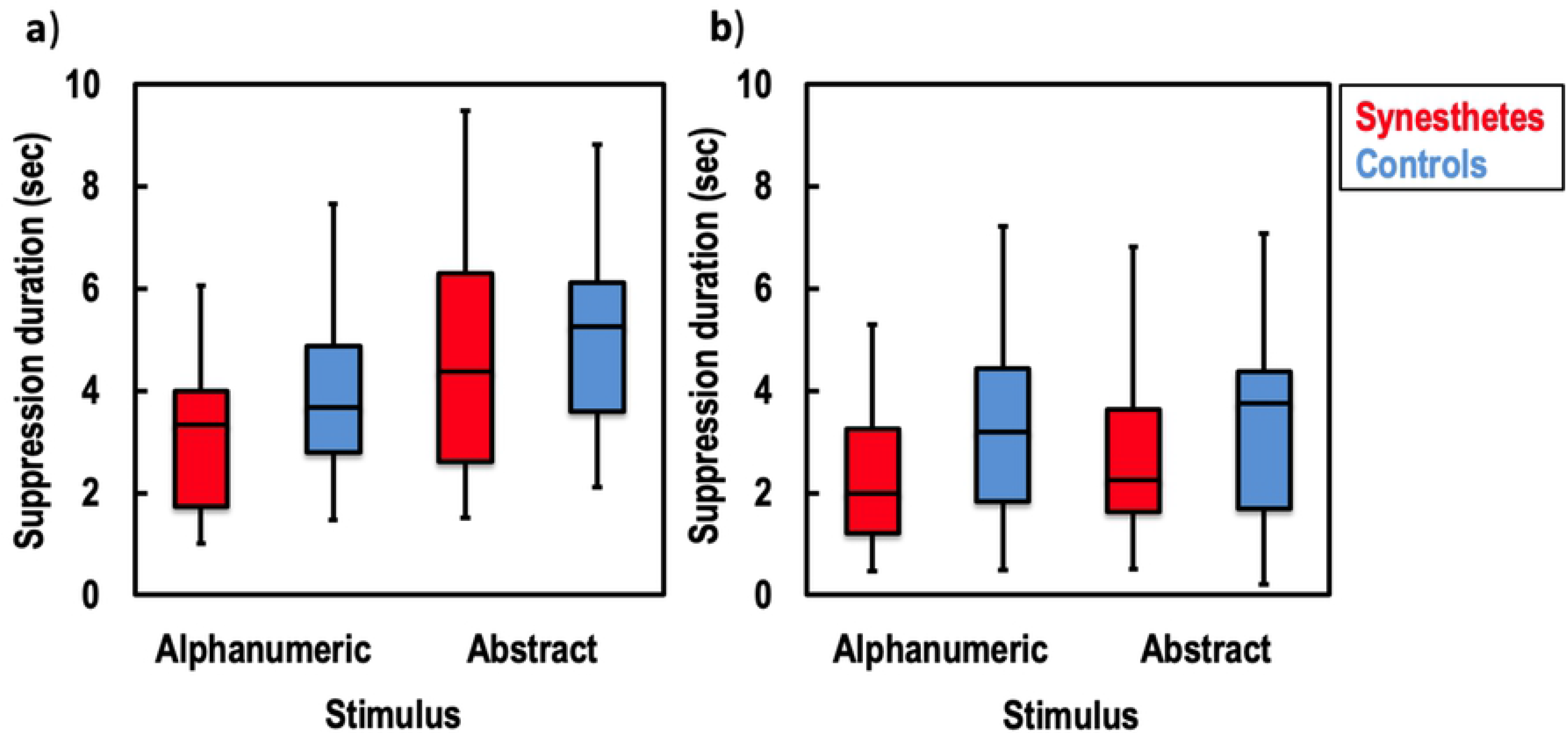
Perceptual stimulus suppression. The flash suppression duration is shown for achromatic (**a**), chromatic (**b**), alphanumeric and abstract stimuli in synesthetic participants (red) and control participants (blue). The whiskers indicate the minimum and maximum range of the distributions; the top and bottom of the boxes show the first and third quartiles (25th and 75th percentile), and the horizontal bars inside the boxes represent the medians.

Two sensitivity analyses were also performed. First, an additional ANOVA was conducted on perceptual suppression duration while excluding the participant reporting a “d” synesthetic stimulus, instead of the “5” observed by the other synesthetes. Second, a similar approach was conducted by excluding the synesthete participant who showed a projector synesthesia profile, as revealed by the online synesthesia battery test. Results from these two ANOVA remained the same (data not shown). It should be noted that *Z*-score tests revealed that the performance of the projector synesthete participant was not significantly different from the other synesthete participants, or from the control participants.

For non-flash-suppression trials, the ANOVA revealed robust significant effects of condition [*F*_(1,19)_ = 21.111, *p* < 0.001, η^2^ = 0.526], stimulus [*F*_(1,19)_ = 23.970, *p* < 0.001, η^2^ = 0.558], and group [*F*_(1,19)_ = 5.651, *p* = 0.028, η^2^ = 0.229]. No interaction effects between factors were found to be statistically significant: condition*stimulus*group [*F*_(1,19)_ = 2.268, *p* = 0.146, η^2^ = 0.107], condition*stimulus [*F*_(1,19)_ = 0.586, *p* = 0.453, η^2^ = 0.030], condition*group [*F*_(1,19)_ = 2.644, *p* = 0.120, η^2^ = 0.122], stimulus*group [*F*_(1,19)_ = 2.877, *p* = 0.106, η^2^ = 0.132]. Thus participants perceived chromatic stimuli faster than achromatic stimuli (Fig 5). In addition, alphanumeric stimuli were more rapidly reported than abstract stimuli. In comparison to controls, target detection in all conditions was in general faster in synesthete participants. Of note, the performance of the synesthetes in the achromatic condition was not statistically different (*t*_(9)_ = 1.231, *p* = 0.249) between the alphanumeric (M= 1.001, SD = 0.164) and abstract stimuli (M = 1.077 SD = 0.216).

**Fig 5.**
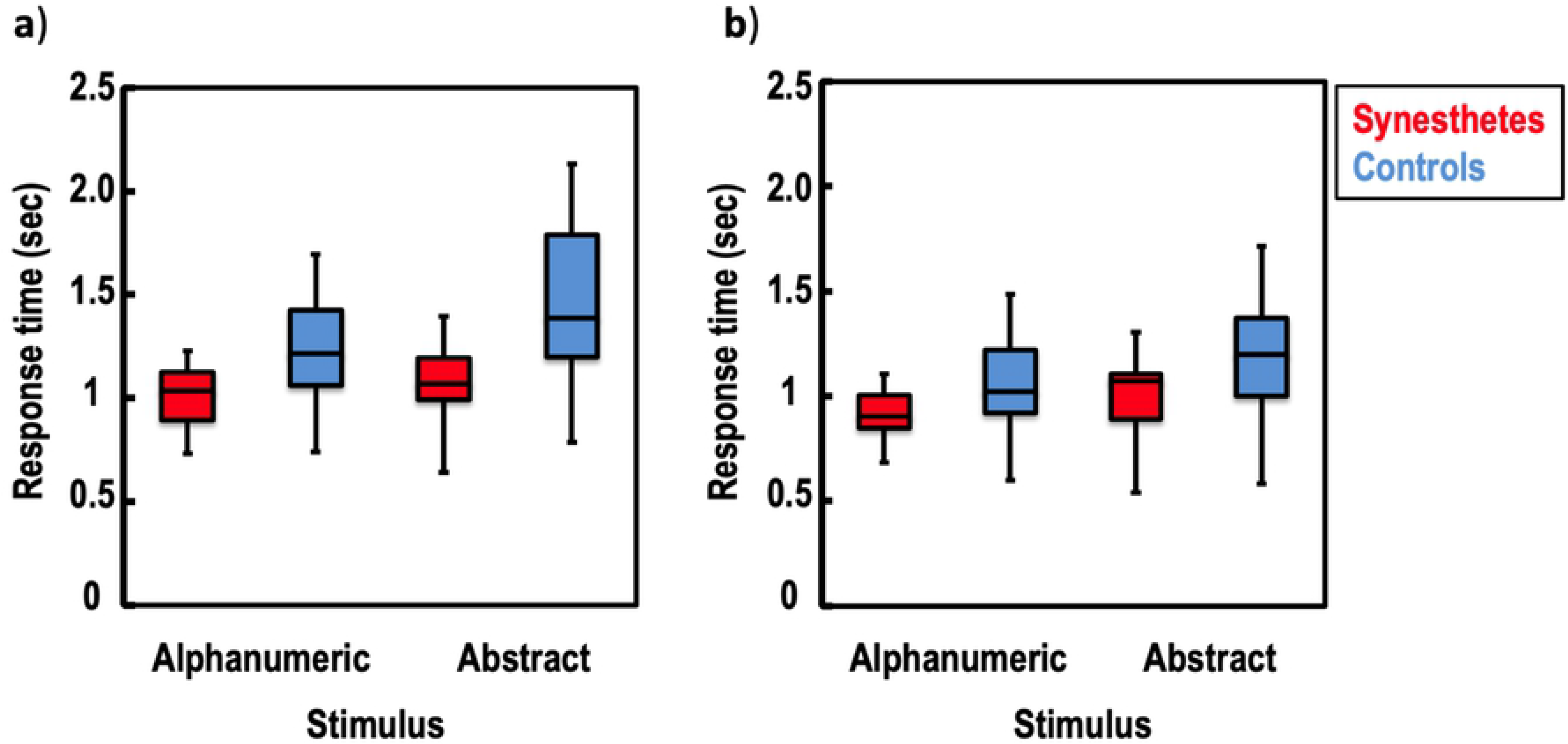
Stimulus detection in absence of flash suppression. Response time is shown for achromatic (**a**) and chromatic (**b**), alphanumeric and abstract stimuli in synesthetic participants (red), and control participants (blue). The whiskers indicate the minimum and maximum range of the distributions, the top and bottom of the boxes show the first and third quartiles (25th and 75th percentile), and the horizontal bars inside the boxes represent the medians.

## Discussion

The present study investigated whether synesthesia shortens the duration of interocular suppression to test the hypothesis that synesthesia may occur in the absence of visual awareness. Results from the flash suppression trials revealed no evidence of suppression modulation from synesthetic stimuli, and no significant differences between synesthete and non-synesthete groups. However, chromatic stimuli exhibited shorter suppression latencies, as they emerge more quickly, than achromatic stimuli. We also found in both synesthetes and controls that, in comparison to the abstract stimuli, the alphanumeric stimuli reduced suppression durations. In the non-flash-suppression trials, all participants detected colored stimuli faster than achromatic stimuli. They also showed shorter reaction times for the detection of alphanumeric stimuli than abstract stimuli. In the non-flash-suppression trials, we found that synesthete participants exhibited a better performance than controls, regardless of the type (synesthetic or not) and the color (chromatic or achromatic) of the stimuli used.

We found that stimulus features, i.e., color and familiarity, influenced the performance in all participants, whether under interocular suppression viewing condition or not. The effect of color on stimulus predominance under dichoptic flash suppression stimulation was previously reported in normal observers [17], which is also in agreement with the notion that color improves signal detection by enhancing the saliency of the stimulus [25, 26]. Regarding the effect of alphanumeric stimuli, it is likely that response time in detection (non-flash-suppression trials), and the duration of the suppression, were shorter because of higher familiarity and/or meaning of those stimuli, in contrast to abstract and unusual symbols. A study conducted by Gobbini and coworkers [27] reported that processing of familiar faces is more likely to resist flash suppression than non-familiar faces. This result is in line with the fact that, under explicit viewing conditions, familiar and semantic stimuli are more efficiently processed and detected than meaningless or unfamiliar targets [for a review see 28]. Our results showed that the effect of stimulus color and familiarity on reducing interocular suppression are not independent, but instead interactive. Thus, colored stimuli overcame suppression faster when they were also familiar.

The current study failed to demonstrate that synesthetic stimuli biased synesthete performance. Indeed, our findings suggest that synesthesia does not bias interocular suppression or modulate stimuli that are rendered temporarily invisible by flash suppression (flash suppression trials). This is in agreement with the masking paradigm study of Mattingley et al. [9], suggesting that conscious recognition is required in order to elicit synesthetic percepts. This interpretation was supported by another study using the attentional blink paradigm, in which no synesthetic effect (i.e., interference stroop-like effect of T2 on subsequent color naming) was observed when the synesthetic prime (T2) was not consciously perceived [13].

The absence of bias by synesthetic stimuli was also observed for the viewing condition exempted from interocular flash suppression. Indeed, response time for synesthetes was not faster for synesthetic stimuli than non-synesthetic in the achromatic condition. Based on the facilitator influence of color in target detection [25], we expected the achromatic synesthetic stimuli to evoke color sensation, as opposed to non-synesthetic stimuli, to improve explicit visual detection. In line with this finding, some studies, under explicit viewing condition, have reported that achromatic synesthetic stimuli do not always show a significant advantage in reaction time over achromatic non-synesthetic symbols [29, 30]. Other studies have failed to find perceptual differences between synesthetes and non-synesthetes. For example, no advantage for synesthetes has been found in some visual search tasks [31, 32], or in the identification of embedded figures [33]. Furthermore, the putative brain atypicalities in synesthetes have been recently challenged [34–36], and many brain mechanisms observed in synesthetes are likely to follow the same rules as those found in non-synesthete individuals [37, 38].

In the explicit detection task (Fig. 5), synesthetes were faster than controls, regardless of whether the stimuli were colored and/or synesthetic. One parsimonious explanation is that synesthetes are better at detecting types of visual stimulus. There is indeed some experimental evidence suggesting that individuals experiencing colored synesthesia show atypical visual processing for non-synesthetic stimuli. For instance, synesthetes show superior color perception compared to controls, not only for hue discrimination [39] but also for luminance and chroma [40]. Moreover, lower contrast discrimination threshold and enhanced performance in color and shape/curvature discrimination tasks have been reported [41, 42]. Other factors may also explain the performance of synesthetes. It has been suggested that synesthetes might exhibit some specific personality traits such as openness or disposition to get involved in new experiences, and that they are even more sensitive to mental imagery [43, 44]. One can reasonably speculate that such subjective particularities might influence perception. Thus synesthetes might differ in how willing they are to affirm the existence of a stimulus, which corresponds with the decision criterion in the signal detection theory, and impacts the observers’ discrimination responses. The faster target detection may therefore be the result of a criterion shift (response bias) rather than a true effect of discrimination. While our study was not designed to verify this possibility, it appears unlikely because the performance of the synesthetes was not different from the controls in the flash suppression condition. In addition, many studies using a signal detection theory design failed to reveal a significant difference in response bias between synesthete and non-synesthete participants [45–47].

Further individual differences in experiencing synesthesia may have influenced the finding of the present study. It is well known that synesthetic experiences differ qualitatively between individuals. For instance, some synesthetes perceive colors as being “outside” of their body, while others perceive them internally, i.e., in the “mind’s eye”. These phenomenological distinctions in synesthetic percepts are known as the projector type and the associator type, respectively [10], although this classification is still under debate [31, 48]. The enhanced performance in synesthesia reported by most studies is best shown for projector synesthetes [10, 24, 49], including for brain activity [50, 51]. For example, the study conducted by Ramachandran et al., [15] suggesting that conscious letter recognition is not required for synesthetic perception, examined only projector synesthetes. By contrast, all synesthetes in the present study were classified as associator types, based on the self-report in the Synesthesia Battery, with the exception of one participant (i.e., ID 2, see Table 1). Of note, the performance of the projector synesthete was not significantly different from the other synesthetes in all tasks. For now, the role of such individual differences in the synesthetic experience in awareness remain poorly understood, as most studies have failed to systematically evaluate and compare participants’ performance with regard to their synesthetic profiles [for example 9, 13–15].

## Conclusion

In summary, our results suggest that synesthesia is less likely to manifest implicitly when stimuli are momentarily invisible; however, a more comprehensive assessment of synesthesia in implicit and mimicked conditions is still necessary to support this interpretation. This could be achieved by designing flash suppression tasks in which individual visual threshold variables, like subject criterion and atypical visual functioning, may be better assessed and controlled [52]; by adapting the flash suppression technique to prime stimuli [i.e., 53]; by including objective measures related to implicit perception as in the recording of neural activity [i.e., 54]; and by also considering individual differences in synesthetic experiences.

## Acknowledgments

We thank Anthony Hosein for his excellent technical assistance.

## References

1. Simner J, Mulvenna C, Sagiv N, Tsakanikos E, Witherby SA, Fraser C, et al. Synaesthesia: The prevalence of atypical cross-modal experiences. Perception. 2006;35(8):1024–33.

2. Dovern A, Fink GR, Fromme ACB, Wohlschläger AM, Weiss PH, Riedl V. Intrinsic network connectivity reflects consistency of synesthetic experiences. Journal of Neuroscience. 2012;32(22):7614–21.

3. Dixon MJ, Smilek D, Cudahy C, Merikle PM. Five plus two equals yellow. Nature. 2000;406(6794):365.

5. Lupiáñez J, Callejas A. Automatic perception and synaesthesia: Evidence from colour and photism naming in a stroop-negative priming task. Cortex. 2006;42(2):204–12.

4. Eagleman DM, Goodale MA. Why color synesthesia involves more than color. Trends in cognitive sciences. 2009;13(7):288–92.

6. Mills CB. Digit synaesthesia: A case study using a Stroop-type test. Cognitive Neuropsychology. 1999;16(2):181–91.

7. Odgaard EC, Flowers JH, Bradman HL. An investigation of the cognitive and perceptual dynamics of a colour–digit synaesthete. Perception. 1999;28(5):651–64.

8. Smilek D, Dixon MJ, Cudahy C, Merikle PM. Synaesthetic Photisms Influence Visual Perception Identification of Masked Digits. 2001:930–6.

9. Mattingley JB, Rich AN, Yelland G, Bradshaw JL. Unconscious priming eliminates automatic binding of colour and alphanumeric form in synaesthesia. Nature. 2001;410(6828):580–2. doi: 10.1038/35069062.

10. Dixon MJ, Smilek D, Merikle PM. Not all synaesthetes are created equal: Projector versus associator synaesthetes. Cognitive, Affective, & Behavioral Neuroscience. 2004;4(3):335–43.

11. Kim C-Y, Blake R. Psychophysical magic: rendering the visible ‘invisible’. Trends in cognitive sciences. 2005;9(8):381–8.

12. Shapiro KL, Raymond JE, Arnell KM. The attentional blink. Trends in cognitive sciences. 1997;1(8):291–6. doi: 10.1016/S1364-6613(97)01094-2.

13. Rich AN, Mattingley JB. Out of sight, out of mind: The attentional blink can eliminate synaesthetic colours. Cognition. 2010;114(3):320–8.

14. Johnson A, Jepma M, de Jong R. Colours sometimes count: awareness and bidirectionality in grapheme-colour synaesthesia. Quarterly journal of experimental psychology (2006). 2007;60(10):1406–22. doi: 10.1080/17470210601063597.

15. Ramachandran VS, Seckel E. Synesthetic colors induced by graphemes that have not been consciously perceived. Neurocase. 2015;21(2):216–9. Epub 2014/03/14. doi: 10.1080/13554794.2014.890728. PubMed PMID: 24621005.

16. Yang E, Brascamp J, Kang M-S, Blake R. On the use of continuous flash suppression for the study of visual processing outside of awareness. Frontiers in psychology. 2014;5:724.

17. Tsuchiya N, Koch C. Continuous flash suppression reduces negative afterimages. Nature neuroscience. 2005;8(8):1096–101. Epub 2005/07/05. doi: 10.1038/nn1500. PubMed PMID: 15995700.

18. Blake R, Logothetis NK. Visual competition. Nature Reviews Neuroscience. 2002;3(1):13.

19. Wolfe JM. Reversing ocular dominance and suppression in a single flash. Vision research. 1984;24(5):471–8.

20. Jiang Y, Costello P, He S. Processing of invisible stimuli: Advantage of upright faces and recognizable words in overcoming interocular suppression. Psychological science. 2007;18(4):349–55.

21. Lupyan G, Ward EJ. Language can boost otherwise unseen objects into visual awareness. Proceedings of the National Academy of Sciences of the United States of America. 2013;110(35):14196–201. Epub 2013/08/14. doi: 10.1073/pnas.1303312110. PubMed PMID: 23940323; PubMed Central PMCID: PMC3761589.

22. Mudrik L, Breska A, Lamy D, Deouell LY. Integration without awareness: expanding the limits of unconscious processing. Psychol Sci. 2011;22(6):764–70. Epub 2011/05/11. doi: 10.1177/0956797611408736. PubMed PMID: 21555524.

23. Hong SW, Blake R. Interocular suppression differentially affects achromatic and chromatic mechanisms. Attention, perception & psychophysics. 2009;71(2):403–11. Epub 2009/03/24. doi: 10.3758/APP.71.2.403. PubMed PMID: 19304629; PubMed Central PMCID: PMC2734383.

24. Eagleman DM, Kagan AD, Nelson SS, Sagaram D, Sarma AK. A standardized test battery for the study of synesthesia. Journal of neuroscience methods. 2007;159(1):139–45.

25. Wolfe JM. Guided search 2.0 a revised model of visual search. Psychonomic bulletin & review. 1994;1(2):202–38.

26. Treisman A. Preattentive processing in vision. Computer vision, graphics, and image processing. 1985;31(2):156–77.

27. Gobbini MI, Gors JD, Halchenko YO, Rogers C, Guntupalli JS, Hughes H, et al. Prioritized Detection of Personally Familiar Faces. PloS one. 2013;8(6):e66620. Epub 2013/06/28. doi: 10.1371/journal.pone.0066620. PubMed PMID: 23805248; PubMed Central PMCID: PMC3689778.

28. Krueger LE. Familiarity effects in visual information processing. Psychological Bulletin. 1975;82(6):949.

29. Sagiv N, Heer J, Robertson L. Does binding of synesthetic color to the evoking grapheme require attention? Cortex. 2006;42(2):232–42.

30. Palmeri TJ, Blake R, Marois R, Flanery MA, Whetsell W. The perceptual reality of synesthetic colors. Proceedings of the National Academy of Sciences. 2002;99(6):4127–31.

31. Edquist J, Rich AN, Brinkman C, Mattingley JB. Do synaesthetic colours act as unique features in visual search? Cortex. 2006;42(2):222–31. doi: 10.1016/S0010-9452(08)70347-2.

32. Nijboer TC, Satris G, Van der Stigchel S. The influence of synesthesia on eye movements: no synesthetic pop-out in an oculomotor target selection task. Consciousness and cognition. 2011;20(4):1193–200. Epub 2011/05/03. doi: 10.1016/j.concog.2011.03.017. PubMed PMID: 21531581.

33. Rothen N, Meier B. Do synesthetes have a general advantage in visual search and episodic memory? A case for group studies. PloS one. 2009;4(4):e5037.

34. Dojat M, Pizzagalli F, Hupe JM. Magnetic resonance imaging does not reveal structural alterations in the brain of grapheme-color synesthetes. PloS one. 2018;13(4):e0194422. Epub 2018/04/05. doi: 10.1371/journal.pone.0194422. PubMed PMID: 29617401; PubMed Central PMCID: PMC5884511.

35. Hupé J-M, Bordier C, Dojat M. The neural bases of grapheme-color synesthesia are not localized in real color-sensitive areas. Cerebral cortex (New York, NY: 1991). 2012;22(7):1622–33. doi: 10.1093/cercor/bhr236.

36. Weiss F, Greenlee MW, Volberg G. No atypical white-matter structures in grapheme-or color-sensitive areas in synesthetes. bioRxiv. 2019:618611.

37. Sagiv N, Ward J. Crossmodal interactions: lessons from synesthesia. Progress in brain research. 2006;155(2001):259–71. doi: 10.1016/S0079-6123(06)55015-0.

38. Arias DJ, Hosein A, Saint-Amour D. Assessing Lateral Interaction in the Synesthetic Visual Brain. Vision. 2019;3(1):7.

39. Banissy MJ, Walsh V, Ward J. Enhanced sensory perception in synaesthesia. Experimental brain research. 2009;196(4):565–71.

40. Banissy MJ, Tester V, Muggleton NG, Janik AB, Davenport A, Franklin A, et al. Synesthesia for color is linked to improved color perception but reduced motion perception. Psychological science. 2013;24(12):2390–7. doi: 10.1177/0956797613492424.

41. Terhune DB, Song SM, Duta MD, Cohen Kadosh R. Probing the neurochemical basis of synaesthesia using psychophysics. Frontiers in human neuroscience. 2014;8(February):89-. doi: 10.3389/fnhum.2014.00089.

42. Ward J, Rothen N, Chang A, Kanai R. The structure of inter-individual differences in visual ability: Evidence from the general population and synaesthesia. Vision Research. 2017; 141:293–302. doi: 10.1016/j.visres.2016.06.009.

43. Chun CA, Hupe JM. Are synesthetes exceptional beyond their synesthetic associations? A systematic comparison of creativity, personality, cognition, and mental imagery in synesthetes and controls. Br J Psychol. 2016;107(3):397–418. Epub 2015/09/09. doi: 10.1111/bjop.12146. PubMed PMID: 26346432; PubMed Central PMCID: PMC5049650.

44. Banissy MJ, Holle H, Cassell J, Annett L, Tsakanikos E, Walsh V, et al. Personality traits in people with synaesthesia: Do synaesthetes have an atypical personality profile? Personality and Individual Differences. 2013;54(7):828–31. doi: 10.1016/j.paid.2012.12.018.

45. Whittingham KM, McDonald JS, Clifford CWG. Synesthetes show normal sound-induced flash fission and fusion illusions. Vision research. 2014;105:1–9.

46. Amsel BD, Kutas M, Coulson S. Projectors, associators, visual imagery, and the time course of visual processing in grapheme-color synesthesia. 2017;(4): 206–223| Cogn Neurosci 8. PubMed Abstract Publisher Full Text.

47. Lunke K, Meier B. New insights into mechanisms of enhanced synaesthetic memory: Benefits are synaesthesia-type-specific. PloS one. 2018;13(9):e0203055.

48. Yokosawa K, Asano M. Relation between synesthetic grapheme-color associations and the sub-types of synesthesia. Journal of vision. 2015;15(12):132-

49. Ward J, Li R, Salih S, Sagiv N. Varieties of grapheme-colour synaesthesia: A new theory of phenomenological and behavioural differences. Consciousness and cognition. 2007;16(4):913–31. doi: 10.1016/j.concog.2006.09.012.

50. van Leeuwen TM, den Ouden HEM, Hagoort P. Effective Connectivity Determines the Nature of Subjective Experience in Grapheme-Color Synesthesia. Journal of Neuroscience. 2011;31(27):9879–84. doi: 10.1523/JNEUROSCI.0569-11.2011.

51. Cohen MX, Weidacker K, Tankink J, Scholte HS, Rouw R. Grapheme-color synesthesia subtypes: Stable individual differences reflected in posterior alphaband oscillations. Cognitive neuroscience. 2015;6(2-3):56.

52. Stein T, Hebart MN, Sterzer P. Breaking Continuous Flash Suppression: A New Measure of Unconscious Processing during Interocular Suppression? Front Hum Neurosci. 2011;5:167. Epub 2011/12/24. doi: 10.3389/fnhum.2011.00167. PubMed PMID: 22194718; PubMed Central PMCID: PMC3243089.

53. Bahrami B, Vetter P, Spolaore E, Pagano S, Butterworth B, Rees G. Unconscious numerical priming despite interocular suppression. Psychol Sci. 2010;21(2):224–33. Epub 2010/04/29. doi: 10.1177/0956797609360664. PubMed PMID: 20424051.

54. Schlossmacher I, Junghofer M, Straube T, Bruchmann M. No differential effects to facial expressions under continuous flash suppression: An event-related potentials? study. NeuroImage. 2017;163:276–85. Epub 2017/09/25. doi: 10.1016/j.neuroimage.2017.09.034. PubMed PMID: 28939431.

